# Adoption of Modern Beekeeping Technologies and Best Management Practices Among Honey Beekeepers in South West, Nigeria

**DOI:** 10.1101/2024.06.16.599159

**Authors:** P.J Adekola, A Aderanti, M. A Oladoja, O Bobadoye Bridget

## Abstract

Colony collapse disorder (CCD) is an abnormal phenomenon which has been largely undocumented in Nigeria. However, several factors have been attributed to high cases of hive abandonment, amongst which is low adoption of hive management practices. This study was carried out to determine adoption of modern technologies and hive management practices amongst beekeepers within the south-west region of Nigeria to reduce the incidence of CCD. A multistage sampling procedure was used to select a total of 399 beekeepers in beekeeping zones across Ekiti, Oyo and Osun and Ondo States in South-West, Nigeria. Interview schedules were used to collect information on beekeepers to determine the level of adoption of modern beekeeping technologies among honey beekeepers to mitigate against colony collapse and results showed that honey beekeepers perception, benefits derived and constraints to adoption of modern honey beekeeping technologies determined what best management practices to adopt. In light of these findings, it is clear that beekeeping in Southwest Nigeria holds significant promise as a source of income and livelihood improvement. To harness this potential, policymakers, government agencies, and relevant stakeholders must consider tailored strategies to address the unique challenges faced by beekeepers in different states. Encouraging the adoption of modern beekeeping technologies and providing the necessary support systems can contribute to the sustainable growth of the beekeeping industry in the region.

In conclusion, there is a need for enhanced education and training initiatives to improve beekeepers’ knowledge of good management practices, which can positively impact beekeeping outcomes. Moreover, addressing the issues related to time constraints and the lack of time for bee farm monitoring is essential to ensure the efficient management of beekeeping activities. The losses and consequent economic damages have encouraged researchers to develop new strategies to control honeybee diseases and pest infestations through the adoption of modern technology.

## INTRODUCTION

Beekeeping is an important activity to agriculture, food security and biodiversity as well as it participates in reducing poverty and boost livelihoods in rural areas worldwide (Chazovachii *et al*., 2013; Gupta *et al*., 2014). It includes collection and care of bee swarms, the pollinated of field crops by bee, the processing of bee products and the breeding of bees for large honey production (Adekola, *el al*, 2009) It also contributes in the agricultural production due to the essential role of honey bees, *Apis mellifera*, in plants pollination as the pollination of more than 90 crops depend mainly on honey bees (Partap *et al*., 2012). About 15 billion dollars have been estimated as pollination value of honey bees in the USA alone in 2000 (Morse and Calderone, 2000). Moreover, honey bees participate in the conservation of the biodiversity for many crops beside their valuable nutritional and medicinal products including; honey, royal jelly and other bee products (Klein *et al*., 2007) which are considered as source of income (Qaiser *et al*., 2013).

Nigeria is blessed with great potential for beekeeping production because of its endowment with favorable environment that comprises fauna and flora resources. It has been part of some traditional agricultural enterprise of many parts in Nigeria, but honey production has always been on the decline and never satisfies local demand (Akinwande, Badejo and Ogbogu, 2013).Traditional beekeeping practice is one of the oldest types of agricultural practices since ages (1000 -1500 AD) years in Nigeria. It is characterized mainly by forest beekeeping that is common in some of the forest covered both in the East, West, North and south Nigeria while the natural colonies of the true honeybees (*Apis mellifera*) build their nests in the hole of tress and under thick branches or in the hollow objects, such as pots, large straws, baskets, e.t.c. especially where there is cool dark sites (Mustsear, 1999). Despite the high potentiality of Nigeria for beekeeping and its extensive practices in different areas across the country, beekeeping research conducted in the nation so far has not managed to characterize and document the apicultural resources and associated constraints of the sector for its proper intervention and utilization to specific potential regions [Chala, *el al*, 2012]. These conditions may generally indicate the importance of considering the biology and ecology of the bees in the study areas and introduce adoption of modern technologies as it were in the developed countries around the world and improve Nigerian beekeeping production. Besides the technological and biological factors, the socio-demographic conditions of beekeepers were observed to play a significant role in the adoption of modern technologies. Thus, it was essential to assess the beekeeping production system as a whole and identify the determinants of hive technology preference and the major constraints of this subsector. According to Adekola, *el al*, (2009) stated that, technology is aimed at improving teaching and learning process in Nigeria as posted in National Policy on Education with overall goal of creating National minds of good conduct of people toward National development at large.. Different case studies and researches are being carried out concerning honey-bee production practices in different areas of the country. However, there is no compiled and tangible information on the honey-bee production practice and hive technology preference of hive technology and management as well as constraints in the potential honey-bee areas in Nigeria..The growth of the global apiculture industry has prompted many beekeepers to adopt modern beekeeping technologies and best management practices in order to maximize honey production, improve hive health, and increase overall beekeeping efficiency. In Nigeria, specifically in the Southwest region, beekeeping has gained significant importance, making it essential for honey beekeepers to adopt these modern techniques and practices. Modern beekeeping technologies and best management practices offer numerous benefits to honey beekeepers, including increased honey production, improved hive health and survival rates, and reduced labor input. Additionally, proper adoption of these practices can also enhance the overall sustainability of the beekeeping industry, ensuring the conservation of honey bee populations and the maintenance of ecological balance. Despite the challenges faced by honey beekeepers in Southwest Nigeria, there have been positive developments in the adoption of modern technologies and best management practices. This can be attributed to the awareness-raising efforts of various government organizations, non-profit entities, and beekeeping associations.

## METHODS AND MATERIALS

### Study Area

The study area is the rain forest of South West Nigeria (Figure 1). South west Nigeria covers Lagos, Ekiti, Ogun, Ondo, Osun and Oyo states which is mainly dominated by the Yoruba ethnic group (Figure 2). This group according to Ayinde (2005) is the largest and the longest established ethnic groups on the African continent. The total population of the area is 27, 581,992 with Lagos state having 9,013,534: Oyo 5,591,589: Ondo 3,441,024; Osun 3,423,535 Ogun 3,728,098 and Ekiti 2,384,212(world population prospect 2006). The area lies in the humid tropical bordered by mangrove swamp forest in the south; major part of the study area lies in the fresh water swamp forest and rain forest to the south which turns to moist and dry woodland savanna towards the north. It has a land area of about 114, 271 square kilometer(12% off the total land mass of Nigeria, lying between Latitude 4’20’ and 9’23’ North Equator and the Longitude 2’25’ and 6’31’ East Agriculture forms the base of the overall development thrust of the zone, with crop farming as the main occupation of the people in the area. Nigeria is characterized by an annual rainfall of about 2000-2500 mm and high humidity. The wet season last between seven and nine months in Ogun, Oyo and Osun and some part of Lagos states fall in the rainforest zone of the south west Nigeria. The soil type in this zone is well drained but leached soils. These element favors the cultivation of crops such as cassava, maize, yam, rice, cocoyam, and beans. In the southwest Nigeria sustainable livestock production is also practiced by agro-pastoralist who are scattered across the zone.

**Figure 1.**
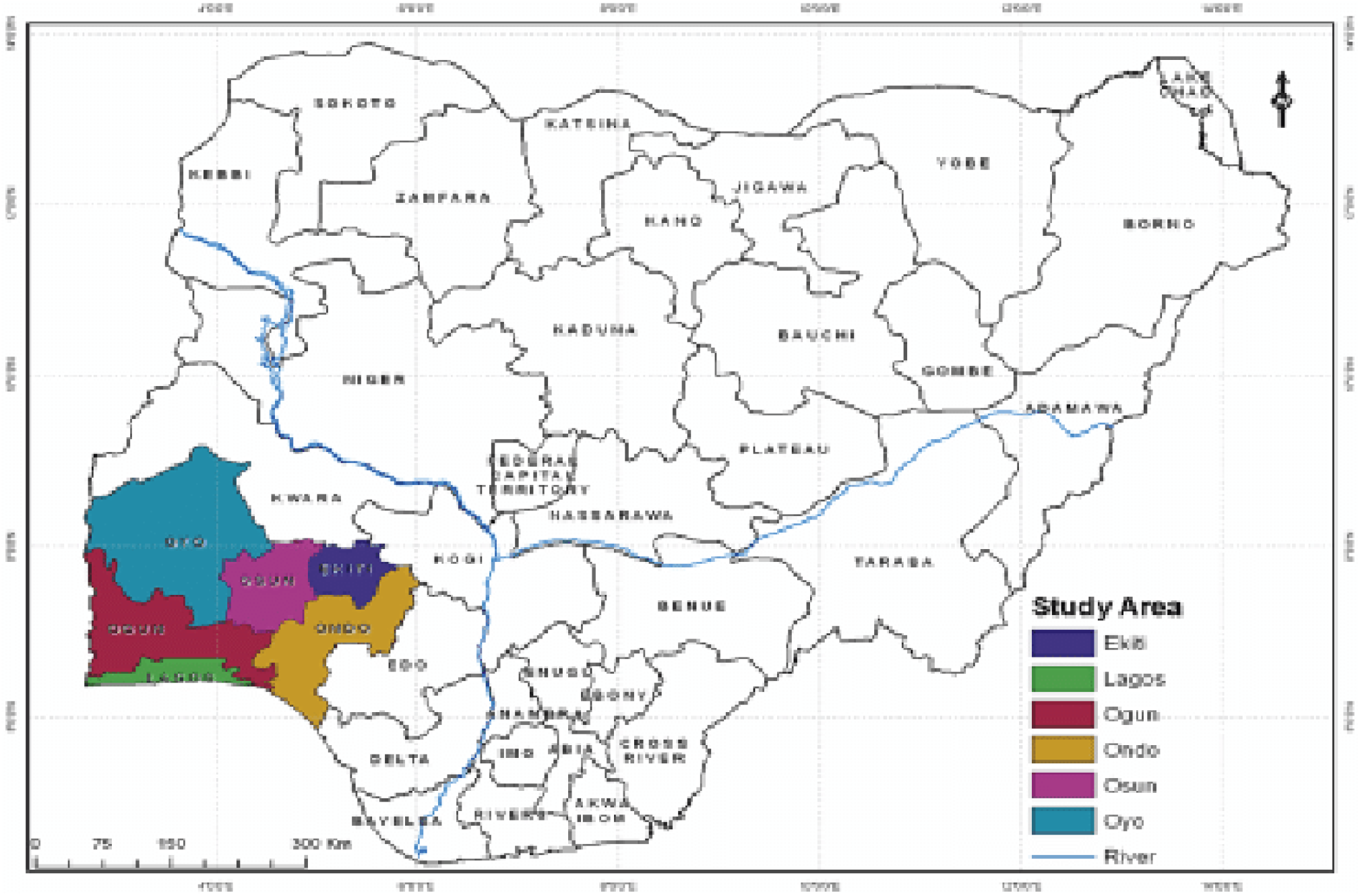
Map of South West Nigeria (Wikipedia, 2023)

**Figure 2:**
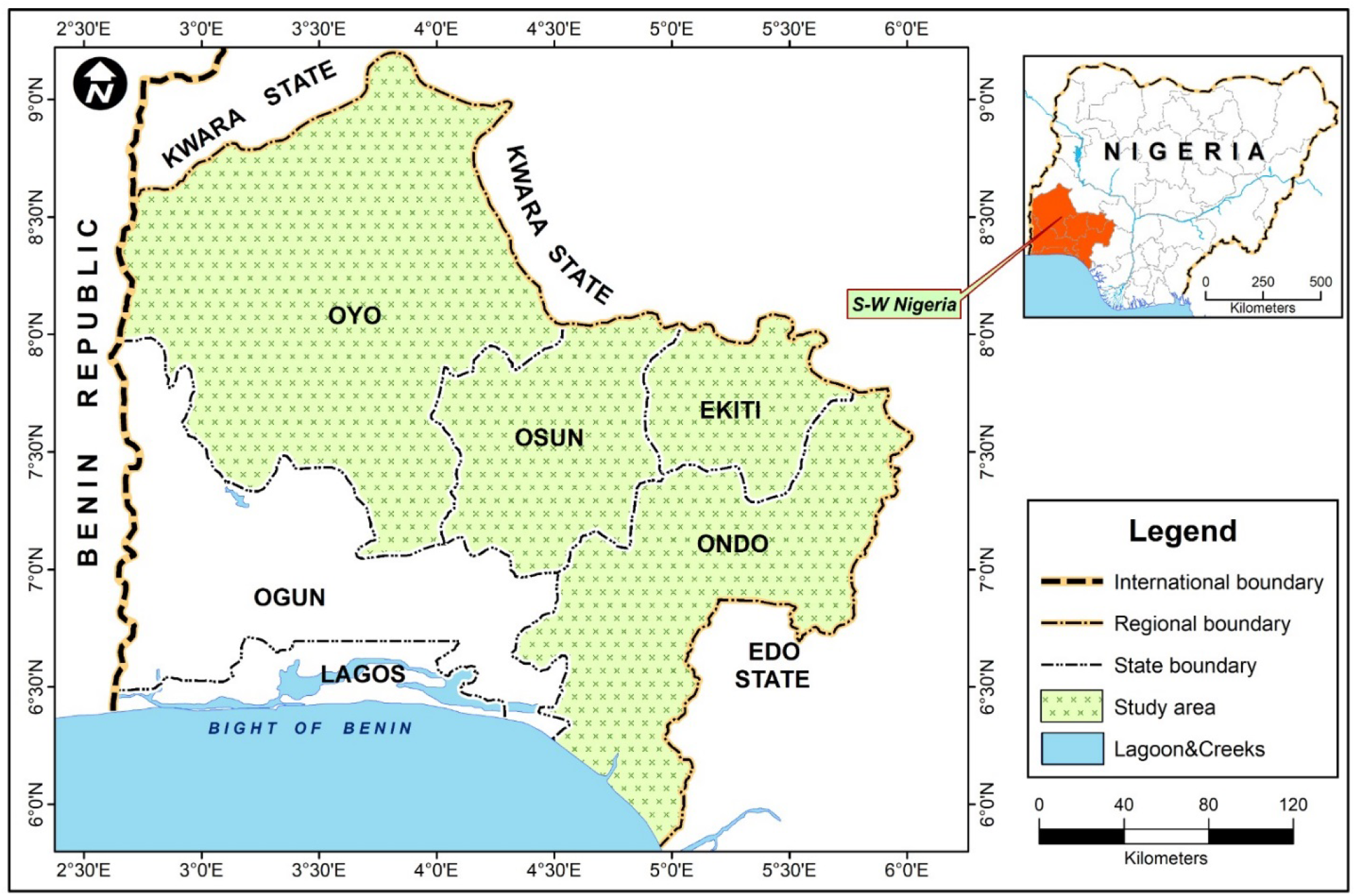
The general objective of the study was to investigate adoption of modern beekeeping technologies among honey beekeepers in southwest, Nigeria.

### Data Collection

Structured interview schedule was used to elicit information from the respondents. Both quantitative and non-quantitative methods were used to collect data from the respondents. Face and content validity of the instruments were conducted by experts in the field of Agricultural Extension. This was done by splitting the questionnaires into equal half, one half composed of even number while other half composed of odd-numbered questions. After administering each half to the same individuals, the correlation between the two halves was calculated after repeating it for the individual. The higher the correlation between the two halves, the higher the internal consistency of the survey.

### Measurement of Variables

Two variables, namely the dependent and independent variables were measured in the study.The independent variables consist of the socio-economic characteristics of the respondents, sources of information, benefits derived from the use of the modern beekeeping technologies and levels of the modern beekeeping technologies. A list of perception statement was presented to the respondent on a 5 points liker scale of SA, A,U,D,SD, The statement covered appropriateness of technology, ease of use of technology, access to technology, affordability of technologies and maintenance of technology.

### Statistical analysis

#### Descriptive Statistics

Descriptive statistics such as frequency percentages, means and standard deviation were used in describing the variables.

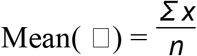

Where Σ x = sum of all the data value, n = number of observation

The standard deviation of ungrouped data values or single scores was calculated using the formula:

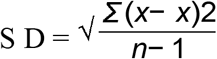

Where S.D = the sample standard deviation,

Σ(*x*−*x*)2 = sum of squares of deviation of the scores from the mean

## RESULTS

### Adoption of beekeeping Technologies

Results in Table 4 indicate extent of adoption of beekeeping technologies both regularly and occasionally in Ekiti states. Kenyan hives are used more than any other equipment like langstroth hives (0-8%) , flow hives (0.0%), bee venom extractor (0-0%), pollen baskets (0.0%) both regularly and occasionally use, queen rearing kits (5.7%), propolis extractor (6.2%), royal jelly extractor (0.0%), lab analysis of bee (0-0%), bee wear/ suits (6%), smokers (64.9%), swarm carcher (89.2%) and others. From the data, it’s evident that Kenyan Hives are notably high, 94.6% to 100.0%. are regularly in Ekiti state than other equipment. This data suggests that Kenyan Hives have gained significant popularity and acceptance among beekeepers in Ekiti state, as they offer advantages that make them the preferred choice for housing honeybees colonies. Alexander *el al*, (2017) affirmed that, Kenyan bee hives last longest and do not require a high degree of knowledge to manage or specialized extraction equipment to collect the honey, on like langstroth hive which is very expensive and require high quality wood attention to detail to maintain bee space.

## Discussion

### Determinants of Modern Beekeeping technology adoption

Our results showed that the dominant factors responsible for the adoption of modern beekeeping technologies for Ekiti, Ondo, Osun and Oyo States respectively were significantly correlated to environmental and anthropogenic factors presented in Tables 1, respectively. Each table presents a descriptive (that is mean and standard deviation, presented as std. dev.) summary for the variables investigated, commonalities (initial and extracted value) as well as summary for the mimportant factors responsible for the adoption. In Table 2, the value of KMO statistic was 0.530 (> 0.50), hence adequate for determining the factors responsible for adopting modern beekeeping technologies for Ekiti State. The significant result (p-value < 0.001) of the Bartlett’s test further establishes that there was relationship between the explanatory variables and adopting modern beekeeping technologies. The study showed factor analysis presented four factors. The communalities column showed the amount of variance in each variable that has been accounted for by the extracted factors. For instance, the variance observed between “Availability of equipment” in Ekiti state was approximately 66.7% to other variables while there was 64.9% variation between item 7 (that is, Affordable cost of equipment) and the other items of interest in this study. On factors responsible for the adoption of modern beekeeping technology in Ekiti state, the availability of equipment was a very crucial factor determining adoption of beekeepers in Ekiti state. This was followed by having “Affordable cost of equipment”, “Customers preference for bee products × “High yield of bee products” and “Access of government supports”.

**Table 1.** Field survey, 2023.

| sn | Variable | Statistics |  | Communalities |  | Factors | Rotated Matrix Factor Score |
| --- | --- | --- | --- | --- | --- | --- | --- |
|  |  | Mean | Std. dev. | Initial | Extraction |  |  |
| 1 | High yield of bee products | 2.53 | 1.621 | 1.000 | **0.745** | F1 | **0.855** |
| 2 | Improved income | 3.37 | 1.617 | 1.000 | **0.812** | F1 | **0.852** |
| 3 | Ease of operation | 4.50 | 1.675 | 1.000 | **0.760** | F1 | **0.811** |
| 4 | Customers preference for the bee products | 5.73 | 1.908 | 1.000 | **0.688** | F2 | **-0.771** |
| 5 | Access to government supports | 5.40 | 2.105 | 1.000 | 0.686 | F1 | -0.583 |
| 6 | Availability of equipments | 4.92 | 2.735 | 1.000 | **0.758** | F2 | **-0.623** |
| 7 | Affordable cost of equipment | 3.37 | 1.640 | 1.000 | **0.822** | F2 | **0.715** |
| 8 | Opportunity for improved expertise | 3.88 | 2.540 | 1.000 | **0.890** | F2 | **0.695** |
| *Kaiser-Meyer-Olkin(KMO) Measure of Sampling adequacy = 0.620;* |  |  |  |  |  |  |  |
| *Bartlett's test (28, 0.05) = 655.146, p-value < 0.001* |  |  |  |  |  |  |  |

**Table 7.** Factors determining adoption of modern beekeeping technology in Ekiti State.

| sn | Variable | Statistics |  | Communalities |  | Factors | Rotated Matrix Factor Score |
| --- | --- | --- | --- | --- | --- | --- | --- |
|  |  | Mean | Std. dev. | Initial | Extraction |  |  |
| 1 | High yield of bee products | 3.27 | 2.232 | 0.309 | **0.594** | F1 | **0.752** |
| 2 | Improved income | 3.30 | 1.808 | 0.306 | **0.436** | F1 | **0.610** |
| 3 | Ease of operation | 3.76 | 1.606 | 0.167 | 0.239 | F2 | 0.458 |
| 4 | Customers preference for the bee products | 4.81 | 2.343 | 0.249 | **0.635** | F4 | **0.781** |
| 5 | Access to government supports | 5.92 | 1.891 | 0.341 | 0.388 | F3 | -0.572 |
| 6 | Availability of equipments | 5.14 | 1.903 | 0.266 | **0.666** | F2 | **-0.802** |
| 7 | Affordable cost of equipment | 5.00 | 2.285 | 0.332 | **0.649** | F3 | **0.786** |
| 8 | Opportunity for improved expertise | 5.38 | 2.240 | 0.351 | 0.402 | F4 | -0.473 |

*Kaiser-Meyer-Olkin(KMO) Measure of Sampling adequacy = 0.530;*
*Bartlett's test (28, 0.05) = 42.712, p-value = 0.037*

### Recommendations

Based on the findings, the following recommendations are hereby proffered:

1. Recognizing the various difficulties faced by beekeepers in the various Southwest Nigerian states is important to providing useful recommendations. Assistant initiatives.
2. should be established which cater for the particular requirements and difficulties faced by beekeepers in each state must be strengthened. States facing problems with hive theft, for instance, may need community awareness campaigns, while states with greater rates of agricultural vandalism may need heightened security measures.
3. Governments at both state and federal levels should consider policies and initiatives that support beekeepers. This includes providing financial incentives, grants, and subsidies to reduce the financial burden of adopting modern beekeeping technologies.
4. Bee keepers should collaborate with other stakeholders for enhanced and broaden agricultural extension services to give beekeepers the direction and assistance they need to adopt better beekeeping management techniques.
5. Start community-wide awareness efforts to address problems like indiscriminate bush burning, which has an impact on beekeepers. Encourage ethical land management techniques that safeguard bee habitats.
6. Government agencies, non-governmental organizations, and agricultural extension services should actively promote the adoption of modern beekeeping technologies among beekeepers in Southwest Nigeria. This includes providing training and workshops on the benefits and proper utilization of modern beekeeping equipments and practices.

## References

Adino A., and Tessema A.,(2021) Determinants of beekeeping adoption by smallholder rural households in Northwest Ethiopia. Journal of Cogent Food & Agriculture homepage: http://www.tandfonline.com/loi/oafa20.

Adekola, J.P.,Fagbenro, J.A, Ojo, M.O., Aderounmu, A.F., Ajibade, Y.A and Olayiwola, I.B., (2009): Technology in Forestry Education and National Development. International Journal of Research Development (NARD).. Vol. 1. No. 2, August, 20009, Pp 367 – 372,

Adejare, S.O. (1990), Beekeeping in Africa. FAO Agricultural Series, Bulletin 68/6. Rome, Italy, pp. 130.

Adekola, P.J., Akoun, J, M. O. Ojo, Fagbeunro, J.A., Aruwayo, A. Ajibade, Y.A. (2009): Approaches to sustainable Apiculture in Nigeria. Journal of forestry Research and Management. December, 2009 Pp88 – 101..

Adekola P..J.. Ogundele. O.J.K., Babatunde, T.O. Ayoola, O.A. (2005) Prospect of Agro-Apiculture As Avertable Tools for Rural and Urban Poverty Reduction, in Nigeria. A Paper Presented At The First National Association of Agricultural Technology ( N.A.A. T.) National Cereals Research Institute (NCRI). Moore Plantation, Ibadan. 23rd – 29th Nov. 2005.

Adekola; P.J. (2019) Life Science Towards Health and Wealth Generation Through Honey Production for Sustainable Development. Continental Journal of Sustainable Development. (2019) 10 (1); 1 – 13. DOI:10.5281/Zenodo.3271174.

Aida, A., Abd-El-Waheed, Hesham.R., El-Seedi (2019) Bee Venom Composition: From Chemistry to Biological activities. Studies in Natural Products Chemistry.

Ajayi, S.I. (1974). The Demand for Money in Nigeria Economy: Comments and Extension. The Journal of Economics and Social Studies.1.

Affognon, H.D.S, Kingori, A.I., Omondi, M.G., Diro, B.W., Murithi S. Makau, S.K. Raaina. (2015) Adoption of modern beekeeping and its impact on honey production in the formalMwingi Distric of Kenya; Assessment Using Honey-Base Impact Evolution Approach Journal of Animal and plant science 24, No 6 .(2015).

